# Comparison of interferometric light microscopy with nanoparticle tracking analysis for the study of extracellular vesicles and bacteriophages

**DOI:** 10.1101/2022.10.07.511248

**Authors:** Romain Sausset, Zuzana Krupova, Eric Guédon, Sandrine Peron, Alice Grangier, Marie-Agnès Petit, Luisa De Sordi, Marianne De Paepe

**Author notes:** E-mail adresses : Romain Sausset Zuzana Krupova Luisa De Sordi Alice Grangier Marie-Agnes Petit Eric Guedon Sandrine Peron.

## Abstract

Research on extracellular vesicles (EVs) and bacteriophages (phages) has been steadily expanding over the past decades as many of their roles in medicine, biology, and ecosystems have been unveiled. Such interest has brought about the need for new tools to quantify and determine the sizes of these biological nanoparticles. A new device based on interferometric light microscopy (ILM), the Videodrop, was recently developed for this purpose. Here, we compared this new device to two nanoparticle tracking analysis (NTA) devices, the NanoSight and the ZetaView, for the analysis of EVs and phages. We used EVs isolated from bacteria, fecal samples, bovine milk and human cells, and phages of various sizes and shape, ranging from 30 to 120 nm of diameter. While NTA instruments correctly enumerated most phages, the Videodrop detected only the largest one, indicating a lower sensitivity threshold compared to the NTA devices. Nevertheless, the performance of the Videodrop compared favorably to that of the NTA devices for the determination of the concentration of eukaryotic EV samples. The NanoSight instrument provided the most precise size distributions but the Videodrop was by far the most time-saving device, making it worthy of consideration for studies conducted on a large number of samples.

## INTRODUCTION

The discovery of the importance of extracellular vesicles (EVs) in basic research, medicine, and industrial applications has placed them in the limelight in the last decades. EVs are produced and released into the environment by eukaryotic and prokaryotic cells, for which they are important actors in intra- and interspecies cell-cell communication. Human EVs are present in numerous bodily fluids, where they participate in numerous homeostatic processes, such as cellular proliferation, and, therefore, in human health and disease (Reviewed in (Shah et al. 2018)). EVs constitute a promising source of biomarkers for many diseases, including cancer and chronic cardiovascular diseases (Lane et al. 2018, Martin-Ventura et al. 2022). Bacterial EVs also play important roles, notably in pathogenesis, as certain bacteria deliver toxic compounds through EVs during infection (Bomberger et al. 2009, Furuta et al. 2009, Bitto et al. 2017, Codemo et al. 2018, Tartaglia et al. 2018, Sahr et al. 2022). Bacterial EVs have also been found to be abundant in aquatic ecosystems (Biller et al. 2014). Finally, EVs have attracted enormous interest in medicine and the pharmaceutical industry as potential vaccines or vectors for the delivery of active therapeutical compounds, as well as in regenerative medicine (reviewed in (Meng et al. 2020) and (Avalos and Forsthoefel 2022)).

The growth in the number of studies and applications involving EVs has driven the development of tools for the characterization of their size and concentration. Such knowledge is important not only for the standardization of studies and procedures (Thery et al. 2018), but also for the use of EVs as biomarkers, as, for example, the concentration and size of circulating EVs has been associated with several cardiovascular diseases (Shah et al. 2018). However, there are still no simple, rapid, and reliable tools to determine the size distribution and concentrations of entire populations of EVs of various size. The large heterogeneity of EV size in most samples, which typically ranges from 20 to 200 nm, even for EVs from a single cell type, requires single-particle measurements to obtain reliable values. In addition, the detection of all EVs requires label-free procedures, as specific molecules or proteins present on the surface of all EVs have not been identified (Thery et al. 2018, Filip 2021). Traditional epifluorescence microscopy (EPI) is not appropriate for EV enumeration and sizing, as EVs are not easily stained with non-specific dyes (i.e., dyes that would stain all EVs), and it does not provide information on the size of objects. On the contrary, transmission electronic microscopy (TEM) can be used to visualize EVs and estimate their approximate size, despite possible underestimation due to shrinkage (Chernyshev et al. 2015, Kotrbova et al. 2019), but is very challenging to use for enumeration.

Therefore, more recent techniques have been developed to determine the concentration and size of nanoparticles, such as tunable resistive pulse sensing (TRPS), nanoparticle flow cytometry (NFCM), dynamic light scattering (DLS), and nanoparticle-tracking analysis (NTA), but these techniques all have certain limitations. TRPS relies on changes in impedance of a nanopore caused by the passage of a nanoparticle in an electrolyte fluid. Although highly valuable and accurate for the characterization of EVs that are relatively homogenous in size (Maas et al. 2014, Doyle and Wang 2019), the dimension of the pores has to be adapted to the size of the analyzed particles, which is not possible for very polydisperse samples (van der Pol et al. 2014). NFCM is a flow cytometry-based technique that has been improved for the detection of nanoparticles (Rossi et al. 2015, Lippe 2018, Zamora and Aguilar 2018), but requires very long observation times or very expensive instruments. DLS is a bulk method that detects the temporal fluctuations of intensities of the light scattered by a population of nanoparticles following illumination by a laser (Doyle and Wang 2019). Although highly valuable for the study of very small particles, as for all bulk methods, it is prone to biases arising from sample heterogeneity (Filipe et al. 2010). NTA is, to date, the most widely used technique for the study of EVs. Like DLS and NFCM, NTA exploits the light scattered by nanoparticles upon illumination with a laser, but the trajectories of single nanoparticles are followed, making it possible to determine their hydrodynamic diameter (Dh), which is the diameter of a hard, perfect sphere with a zero surface charge that would diffuse at the same speed as the measured particle (for simplicity, we will use size to indicate Dh when discussing measurements provided by NTA). The ZetaView (Particle Metrix, Germany) and NanoSight (Malvern, UK) are two devices that rely on NTA commonly used in the EV field. Although they are based on the same principle, the ZetaView and NanoSight present important differences in the composition of their hardware and software (Bachurski et al. 2019). For example, with the NanoSight instrument, a flux is generally applied to the sample during acquisition, whereas the acquisition is recorded on a static sample with the ZetaView. In addition, ZetaView is capable of measuring particle motion under an applied electric field, which allows calculation of the zeta-potential, a proxy for particle surface charge. On polystyrene and silica nanospheres, the ZetaView was shown to provide better concentration measurements, whereas the NanoSight provided greater precision in size estimations (Bachurski et al. 2019).

Despite the strengths of NTA, its main limitation results from the sixth-power dependence of the scattered light intensity on the size of the particle, resulting in inaccurate measurements in polydisperse samples (Gardiner et al. 2013, Dehghani et al. 2021). In addition, the use of NTA-based devices is relatively time-consuming (approximately 10 and 20 minutes per sample for the ZetaView and the NanoSight, respectively), which is a practical limitation for studies that require a large number of samples. Therefore, there is still a need for a tool that is both very rapid (i.e., acquisition of results in less than a few minutes) and appropriate for highly polydisperse samples. These properties are the advertised strengths of a new instrument based on interferometric light microscopy (ILM), the Videodrop (Myriade, France). As NTA, ILM uses Brownian motion to calculate the size distribution of the analyzed particles. However, contrary to NTA, a simple LED illuminates the samples and a transmission bright-field microscope is used as a homodyne interferometer to detect the particles due to the interference created by the superposition of the incoming light field and the light scattered by the nanoparticles. Being mostly in the Rayleigh scattering regime, the intensity of the scattered amplitude is proportional to the third-power of the particle diameter (Hulst 1957) and not to the sixth-power, as in NTA, limiting the decrease of signal intensity with particle size.

The Videodrop was developed from an interferometer (Boccara et al. 2016, Roose-Amsaleg et al. 2017), which has been used to enumerate nanoparticles in cheeses (Dugat-Bony et al. 2020) and aquatic environments (Boccara et al. 2016, Roose-Amsaleg et al. 2017). A recent study showed the Videodrop to correctly enumerate two lentiviral virions and a baculovirus, whereas it underestimated the absolute viral concentration of an adenovirus relative to classical titration procedures (Turkki et al. 2021). A Videodrop instrument was also used to estimate the concentration of EVs in plasma, but no control experiments to verify the performance of the device were done in this study (Sabbagh et al. 2021). The main characteristics of the Videodrop, ZetaView, and NanoSight instruments are summarized in Table 1.

**Table 1.** General characteristics of the Nanosight, the ZetaView, and the Videodrop devices. Information was collated using manufacturer websites.

|  | Videodrop | ZetaView | NanoSight |
| --- | --- | --- | --- |
| Technology | ILM | NTA | NTA |
| Light source | LED | Laser | Laser |
| Sample introduction | Droplet in a tank | Syringe, tubings | Syringe, tubings |
| Acquisition time | 20 sec | 1 min | 1 min |
| Temperature control | No | Yes | Yes |
| Possibility of Fluorescence | No | Yes | Yes |
| Cost of consumables | ~ 0 | \$ | \$ |
| Cost of equipment | \$ | \$\$ | \$\$ |

In this study, we compared the performance of the Videodrop in determining the size and concentration of EVs with that of two NTA devices, the ZetaView and the NanoSight. To strengthen our analysis, we expanded our methodological comparison to biological nanoparticles of similar sizes but of different nature, bacteriophages (or simply phages). The phage particles represented benchmark comparisons, as they constitute monodisperse populations, the diameter of the capsid being highly homogenous among virions of the same species. Importantly, the refractive indices (on which depend the signal intensity) of viruses are relatively similar to those of EVs (between 1.42 and 1.49 for viruses versus 1.36 to 1.39 for EVs) (Holzwarth et al. 1974, Chandler et al. 2011, van der Pol et al. 2012, Gardiner et al. 2013, Pang et al. 2016). In addition, in contrast to EVs, phages can be reliably quantified using various techniques, such as plaque counting, quantitative PCR and EPI. Furthermore, phage enumeration is of great interest in itself, since phages, as predators of bacteria, are important actors in all microbial ecosystems and, notably, in human-associated microbiota (Sausset et al. 2020). Differences in phage composition have been shown, for example, to be associated with intestinal bowel diseases (IBDs), such as Crohn’s disease and ulcerative colitis (Norman et al. 2015, Cornuault et al. 2018, Clooney et al. 2019). The rapid and reliable enumeration of phage particles is therefore of considerable interest, not only for understanding the dynamics of complex microbial ecosystems, but also for the development of phage-based applications in biotechnology and medicine.

Here, we present the performance of the ILM and NTA devices for the observation of purified EVs and phages. We used nine types of EVs of very different origin, i.e., originating either from milk, human or bacterial cells, or rodent feces (germ-free or raised conventionally), and purified using a number of different procedures (differential centrifugation, iodixanol or sucrose density gradient and size exclusion chromatography). The phages were chosen to cover a large range of capsid diameters, from 30 to 120 nm, and to have different morphotypes: we used myophages and siphophages, which have long protein tails, and podophages and *Tectiviridae*, which have no or very short tails (**Table 2**). We also compared the size distributions obtained using the Videodrop and NTA devices to those obtained by transmission electron microscopy (TEM), knowing that the sizes derived from Brownian motion do not equate with the geometrical diameters given by TEM and that the size of EVs can be reduced by 15 to 30% during TEM observations (Chernyshev et al. 2015).

**Table 2.** Characteristics of the phages used in this study. Capsid diameters are derived from Viral Zone (viralzone.expasy.org), except for SPP1 (Ignatiou et al. 2019).

|  | T4 | T5 | λ | SPP1 | ΦcrAss001 | T7 | PRD1 | ΦX174 |
| --- | --- | --- | --- | --- | --- | --- | --- | --- |
| Family or morphotype | Myophage | Siphophage | Siphophage | Siphophage | Podophage | Podophage | *Tectiviridae* | *Microviridae* |
| Capsid diameter (nm) | 120 x 86 | 90 | 60 | 61 | 76 | 60 | 66 | 30 |
| Genome type | dsDNA | dsDNA | dsDNA | dsDNA | dsDNA | dsDNA | dsDNA | ssDNA |
| Genome size (kb) | 169 | 121 | 48 | 44 | 103 | 40 | 15 | 5 |

We show that the Videodrop is less sensitive than the NTA devices but that it estimates concentrations similar to those obtained by NTA for most EVs derived from mammalian cells. In addition, the Videodrop was by far the most time-saving device, making it the most appropriate for studies conducted on a large number of samples comprised of sufficiently large objects.

## MATERIALS AND METHODS

### Phage lysates

*Escherichia coli* and *Bacillus subtilis* cultures were grown at 37°C in Luria broth (LB) supplemented with 5 mM MgSO_4_, 5 mM CaCl_2_, and 0.2% maltose for lambda phage. The cultures were infected during exponential growth (OD_600 nm_ of 0.2). *E. coli* phages were grown on the MG1655 *hsdRM* strain (MAC 1403, kanR) at a multiplicity of infection (MOI) between 0.1 and 0.25, except for PRD1, which was grown at 37°C on *E. coli* HMS174 pL2 (with kanamycin at a final concentration of 25 μg/mL), and ΦX174, which was grown at 30°C on *E. coli* CQ2. For both a MOI of 5 was used. SPP1 was grown on *B. subtilis* YB886 using a MOI of 0.01. ΦCrAss001 was grown as described by Shkoporov *et al*. (Shkoporov et al. 2018). Briefly, *Bacteroides intestinalis* (DSM 108646) was grown in fastidious anaerobic broth (FAB) under anaerobic conditions and infected at a MOI of 1. All infections were performed in 500 mL exponentially growing bacterial cultures (OD_600 nm_ of 0.2) and incubated until deceleration of bacterial growth was observed. Bacteria and debris were then pelleted by centrifugation at 5,200 x g for 30 min and the supernatants filtered using a vacuum-driven Stericup filtration system at 0.22 μm (Merck Millipore).

### Extracellular vesicle preparations

Bacterial culture supernatants containing EVs were obtained from 500-mL cultures grown at 37°C. *Faecalibacterium prausnitzii* L2-6 was grown in an anaerobic chamber filled with 5% H_2_, 5% CO_2_, and 90% N_2_ in sterile brain heart infusion supplemented (BHIS) medium supplemented with L-cysteine (0.5 g/mL), maltose (1 g/mL), and cellobiose (1 g/mL) for 24 h. *Staphylococcus aureus* HG003 was grown in BHIS medium and *B. subtilis* YB886 and *E. coli* Nissle 1917 in LB, all with 150 rpm/min agitation in a 1 L flask for 18 h. All cultures were centrifuged at 10,000 x g for 20 min and the supernatant filtered through 0.22-μm Stericup Millipore filters. EVs from murine feces and cecal content were collected either from eight-week-old C57BL/6NRj males (Janvier Labs) or 10-week-old germ-free C3H/HeN and C57BL/6J mice from the Anaxem animal facility (INRAe, Jouy-en-Josas, France), both maintained in a 12 h-light/12 h-dark cycle and fed a chow diet (Ssniff). Rat feces was collected from germ-free F344 Fisher rats grown in the Anaxem facility. To obtain fecal filtrates, fresh or frozen (at −80°C, immediately after sampling) material was diluted 40-fold in cold 10 mM Tris (pH 7.5) and resuspended by gentle agitation at 4°C for 15 min. After centrifugation at 5,200 x g for 30 min, the supernatant was filtered using 0.22 μm pore-size Pall Acrodisc syringe filters.

### Purification of bacterial EVs and phages

Phage lysates and EVs from *B. subtilis* and *F. prausnitzii* supernatants, prepared as described above, were concentrated by centrifugation at 20,000 x g for 16 h (rotor SS-34, Sorvall RC 5C PLUS). Pellets containing phage were resuspended in 0.5 mL SM buffer (200 mM NaCl, 50 mM Tris pH7, 10 mM MgSO_4_) and those containing EVs in Tris 10 mM pH7. Resuspended pellets were then subjected to an iodixanol density gradient in 5-mL Ultra-Clear centrifuge tubes (Beckman Coulter). The tubes were first filled with a two-layer gradient, in which 1.875 mL 45% iodixanol was gently injected under 2.5 mL 20% iodixanol, and the resuspended pellets then layered on the top. After ultracentrifugation at 200,000 x g for 5 h at 4°C (SW55Ti rotor in a XL-90 Beckman Coulter centrifuge), fractions containing purified EVs or purified phages were extracted from the top following the scheme in **Supp. Fig. 1**. Fractions of interest were dialyzed overnight at 4°C against 1 liter of 10 mM Tris for EVs or SM buffer for phages, under agitation, using 25 kD Spectra/Por dialysis membranes. A second dialysis of 3 h was performed the next day under the same conditions. All samples were stored at 4°C and analyzed within the following two weeks. TEM was performed to verify the quality of all EV samples. Cell-free supernatants from *S. aureus* cultures were subjected to EV isolation and purification by sucrose density gradient ultracentrifugation, as described previously (Luz et al. 2021). Cell-free supernatants from *E. coli* cultures were subjected to EV isolation and purification by size-exclusion chromatography, as described previously (Rodovalho et al. 2020).

### Enrichment of THP1 EVs

THP1 cells (ATCC) were cultivated in RPMI medium supplemented with 10% FBS and 1% penicillin-streptomycin at 37°C in 5% CO_2_. They were maintained at a concentration of between 3 × 10^5^ and 1 × 10^6^ cells/mL. For EV production, the cells were washed with RPMI + 1% PS without serum and then resuspended in the same medium in T175 flasks at a concentration of 2.5 × 10^5^ cells/mL in 50 mL. After 48 h, the conditioned media was harvested and centrifuged for 10 min at 2,000 x g. The supernatant was ultracentrifuged for 2 h at 150,000 x g in an Optima XP centrifuge (Beckman) with a MLA-50 rotor. The EV pellet was resuspended in 0.8 mL sterile PBS. The EVs were aliquoted and stored at −80°C until further analysis.

### Isolation of bovine milk-derived EVs

Whole bovine milk samples (200 mL) were centrifuged at 3,000 x g for 15 min at 4°C (Allegra X-15R, Beckman Coulter, France) to separate the fat from the skimmed milk. The whey was obtained after acid precipitation of the skimmed milk with 10% (v/v) acetic acid at 37 °C for 10 min followed by the addition of 10% (v/v) 1 M sodium acetate and a further incubation of 10 min at room temperature. The precipitate was then centrifuged at 1,500 x g at 4°C for 15 min. The supernatant was filtered using a vacuum-driven 0.22-μm filtration system Steritop (Merck Millipore). The whey supernatants were concentrated by centrifugation at 4,000 x g at 20°C using Amicon 100-kDa centrifugal filter units (Merck Millipore) to a final volume of ∼6 mL. Aliquots of 500 μl of the obtained retentate were loaded onto a qEVoriginal 70 nm SEC column (Izon Science, New Zealand) previously washed and equilibrated with PBS. Fraction collection (0.5 mL per fraction) was immediately carried out using PBS as the elution buffer. The selected elution fractions (1-3 of 500 μL each) were pooled and subsequently concentrated using 100-kDa Amicon centrifugal filter units (Merck Millipore). The concentrated samples were subjected to several washing steps with PBS to obtain a highly pure EV population. The EV standards were aliquoted and stored at −80°C until further analysis.

### Transmission electron microscopy (TEM)

Purified EVs or phage samples (10 μL) were directly adsorbed onto a carbon film membrane on a 300-mesh copper grid, stained with 1% uranyl acetate dissolved in distilled water, and dried at room temperature. Grids were examined using a Hitachi HT7700 electron microscope operated at 80 kV (Elexience) and the images acquired with a charge coupled device camera (AMT). This work was carried out at, and with the expertise of, the MIMA2 platform, INRAE (Jouy-en-Josas, France).

### Plaque assay (PA)

Ten microliters of the appropriate dilution of the purified phage preparations were mixed with 300 μL of a culture of their bacterial hosts grown overnight in the conditions used for phage lysates (see above). For all phages, except ΦcrAss001, the phage-bacteria suspensions were mixed with 5 mL warm soft top agar (0.45% w/v agar, 0.25% w/v NaCl, 0.1% w/v Bacto Tryptone; in osmosis-purified water) and immediately poured into Petri dishes already containing a solid LB agar layer (1.5% w/v agar and 2.5% w/v LB powder) in triplicate. For ΦcrAss001, the published protocol was followed (Shkoporov 2018). Briefly, the anaerobically prepared phage-bacteria suspension was mixed in 5 mL 0.4% Bacto agar and immediately poured into three Petri dishes already containing a solid layer of fastidious anaerobic agar (FAA). After solidification, the Petri dishes were incubated overnight at 37°C, except for ΦX174 (30°C). The next morning, lysis plaques were manually counted and the phage titers in plaque-forming units per mL (PFU/mL) were calculated.

### Quantitative PCR (qPCR)

DNA standards for calibration were prepared from phage genomic DNA as follow. Prior to genomic extraction, 500 μL of high-titer phage samples were treated with 0.50 μL Turbo DNAse I (Ambion, 2 U/μL) and 1 μL RNAse A (10 mg/250 mL) at 37°C for 30 min. After adding EDTA to a final concentration of 10 mM to inactivate nucleases, phage DNA was extracted by two phenol-chloroform-isoamyl alcohol (25:24:1) extractions followed by three chloroform-isoamyl alcohol (24:1) purification steps. DNA was precipitated with two volumes of ethanol and 300 mM potassium acetate pH 4.8 and resuspended in 10 mM Tris buffer pH 8. DNA was quantified using a Qubit® device and diluted in 10 mM Tris pH 8 to obtain a concentration 5.0 × 10^6^ genomes of phage in 6 μL. The sample was further diluted four times 3 fold to obtain the calibration range. Phage samples were quantified after treatment with Turbo DNAse I (1 μL for 1 ml of phage sample diluted 10 times in Tris 10mM pH8) for 1 h at 37°C followed by 30 minutes at 95°C to explode capsids and degrade the DNase. The primers shown in **Supp Table 1** were used at a concentration of 10 μM. qPCR was performed in a total volume of 15 μL in MicroAmp Fast Optical 96-well plates sealed with MicroAmp Optical Adhesive Film using the Takyon ROX SYBR Mastermix blue dTTP kit. Amplifications were run in duplicate on a StepOnePlus real-time PCR system with the following cycling conditions: 95°C for 5 min, (95°C for 15 s, 58°C for 45 s, 72°C for 30 s) for 45 cycles, 72°C for 5 min, 95°C for 15 s, 60°C for 15 s, 95°C for 15 s. All phage preparations were independently quantified three times. The analysis of the melting curves confirmed the specificity of the primers. Data analysis was performed using the manufacturer’s StepOne Software 2.3.

### Epifluorescence microscopy (EPI)

Phage samples were diluted to a concentration of approximately 10^7^ virions per mL. Glutaraldehyde was added to 1 ml of diluted sample to a final concentration of 0.5% and incubated at 4°C for 15 min. Samples were then flash frozen in liquid nitrogen. After thawing, 4 mL SM buffer was added to each sample prior to filtration on 0.02-μm Anodisc filters. Each filter was then incubated on a 50-μL drop of SYBR Gold at 200X in the dark for 15 min, with the virus side up. After removal from the drop, filters were dried in the dark before being mounted on a glass slide with Fluoromount-G and a coverslip. Slides were stored at −20°C until observation. Microscopic observations were carried out using a Nikon Ti-E fitted with a 100× oil objective Apo TIRF (NA, 1.49; Nikon) with an iLas2 laser coupling system from Gataca Systems (150 mW, 488 nm). Ten images were captured per slide in the bright field and GFP fluorescence channels (with an excitation filter wavelength of 472/30 nm and emission filter wavelength of 520/35 nm). Emission was collected using interference filters and the images captured using a pair of sCMOS cameras (Orca Flash 4.0 v2 sCMOS; Hamamatsu), with the gain defined at 300, attached to a ×2.5 magnification lens, with a time exposure of 100 to 200 ms, depending on the fluorescence intensity of the phage. The final pixel size was 64 nm. Metamorph v.7 software packages were used to control and process the image acquisition and the images were further analyzed using ImageJ (v1.52a). The number of phages on the whole filters was calculated by multiplying the average counts by the quotient of the area of the filter in contact with the phages by the area of the images.

### ILM measurements with the Videodrop

All purified samples were diluted to appropriate concentrations (20-100 particles per frame) in their respective buffers and 6-μL drops were used for the measurements. Triplicates of the accumulation for 10 acquisitions of 100 frames were recorded per sample in accumulation mode. In accordance with the manufacturer’s specifications, a relative threshold of 3.8 was applied for detection. The removal of macroparticles was enabled using a minimum radius of 10 and a minimum number of hot pixels of 80, as well as drift compensation. Concerning the tracking settings, a maximum of two jumps was tolerated for a minimal length track of 10 frames. The doublet detector (qvir software version 2.5.2.6196) was used for all measurements. Size distributions were obtained after the application of a mobile mean with a period of 3 on histograms with classes of 5 nm. For linearity measurements, polystyrene beads (Polystyrene Nanosphere Suspension Series 3000, ThermoFisher) of various sizes were diluted in water after a short sonication. Bead concentrations were calculated following the indications of the Nanosphere supplier provided in the technical guide.

### NTA measurements with the PMX 220 ZetaView

The Zetaview system (Particle Metrix, Germany) was equipped with a 488-nm laser. Measurement concentrations were obtained by pre-diluting the samples to the ideal 50–200 particles/frame. Each experiment was performed in duplicate on 11 different positions within the sample cell with the following specifications and analysis parameters: cell temperature 25°C, sensitivity 70, shutter 100, Max Area 1,000, Min Area 10, and Min Brightness 25. The results were validated while obtaining at least 1,000 valid tracks for each run. For data capture and analysis, Nanoparticle Tracking Analysis Software (ZNTA) v 8.05.04 was used. Size distributions were obtained after the application of a mobile mean with a period of 3 on histograms with 5-nm classes.

### NTA measurements with the NanoSight NS300

All purified samples were diluted in their appropriate sterile buffer to a volume of 1 mL at an ideal concentration of 20-100 particles per frame and injected at a speed of 50 μL/s into the machine’s specimen chamber with a 1-mL sterile syringe. For each measurement, five acquisitions of 1 min were recorded at 25°C.

The device was equipped with a sCMOS camera and a laser module of 488 nm for most experiments, except for phages PRD1 and ΦX174 and milk EVs, for which a laser module of 405 nm was used. Observations of T4 and PRD1 phages with both laser modules (405 nm and 488 nm) showed almost no differences in concentration or size. A camera level of 15 was used in all experiments. For each measurement, five 1-min videos were captured. After capture, the videos were analyzed using NanoSight software NTA 3.3 Dev Build 3.3.104, with a detection threshold of 4. Size distributions were obtained after the application of a mobile mean with a period of 3 on histograms with 5-nm classes using Excel.

## RESULTS

### Evaluation of the size threshold and linearity range of the Videodrop on polystyrene beads

First, we observed standardized polystyrene beads of known sizes and concentrations at various dilutions with the Videodrop. There was good agreement between the theoretical and measured concentrations between 5 × 10^8^ and 4 × 10^9^ beads/mL for beads with a diameter > 80 nm (**Fig. S2**). The concentrations of 70-nm beads was underestimated by 5 to 10-fold, indicating that 80 nm is the Videodrop size threshold for the good enumeration of polystyrene beads, in agreement with the manufacturer’s indications (**Table 1**). By comparison, the NanoSight was previously shown to provide good agreement between theoretical and measured concentrations for 100-nm diameter polystyrene beads at concentrations between 2 × 10^7^ and 3 × 10^9^ particles/mL (Maguire et al. 2017).

### Determination of phage concentration

We then evaluated the performance of the Videodrop and the two NTA devices for measuring the concentrations and sizes of our benchmark biological nanoparticles, i.e., phages of known concentrations and sizes. Thus, we first purified phage virions of eight species belonging to different taxonomic families with different virion shapes and sizes (**Table 2**). TEM observations showed the phage preparations to be almost free of EVs and other large contaminants (**Fig. 1**). When non-phage objects were observed, as in the PRD1 and T5 preparations, they were more than 10-fold less abundant than the phage particles. In the PRD1 and ΦX174 preparations, the virions appeared to be aggregated on the TEM images, but the subsequent size measurements obtained by NTA indicated that this aggregation occurred on the TEM grids. The capsid and tail dimensions measured for each phage were close to published values (**Tables 2 & 3**).

**Figure 1.**
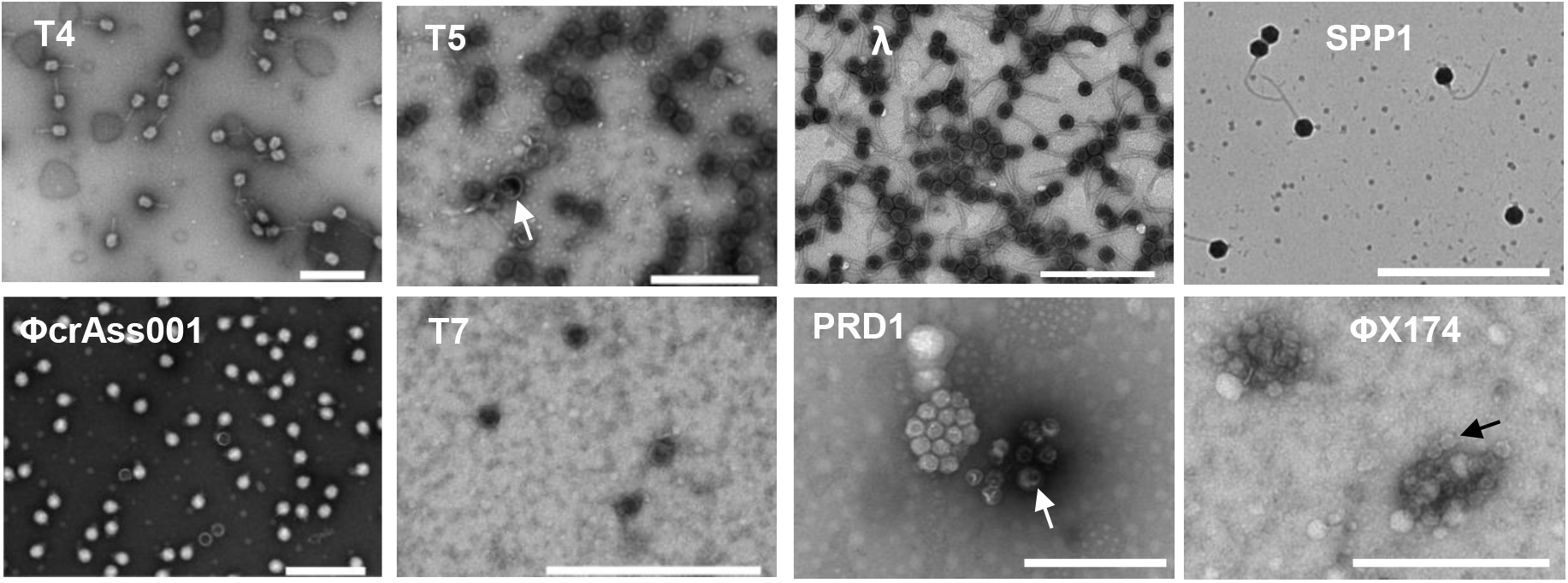
Quality control of phages used for the comparison of the Videodrop, the ZetaView and the NanoSight. Representative TEM images of phage samples. White arrows point to EVs in the PRD1 and T5 stocks. The black arrow in PhiX174 points to a virion. Scale bars are 500 nm.

The concentrations of most phage preparations were determined using three reference methods: EPI, PA, and qPCR. EPI and qPCR provided comparable concentrations, comforting the reliability of the techniques (**Fig. 2**). We thus used the concentrations obtained by EPI as the reference value for further comparisons. The concentrations obtained by PA were only slightly lower than those obtained by EPI and qPCR for most of the phage samples (2 to 3-fold lower), except for T7 and SPP1, for which the differences were greater. This is likely related to the presence of non-infectious virions, as regularly reported (Huang and Baltimore 1970, Heider and Metzner 2014), in particular after thorough purification, which can damage the particles. In support of this hypothesis, we observed tailless phage particles in the SPP1 stocks (**Fig. 1**).

**Figure 2.**
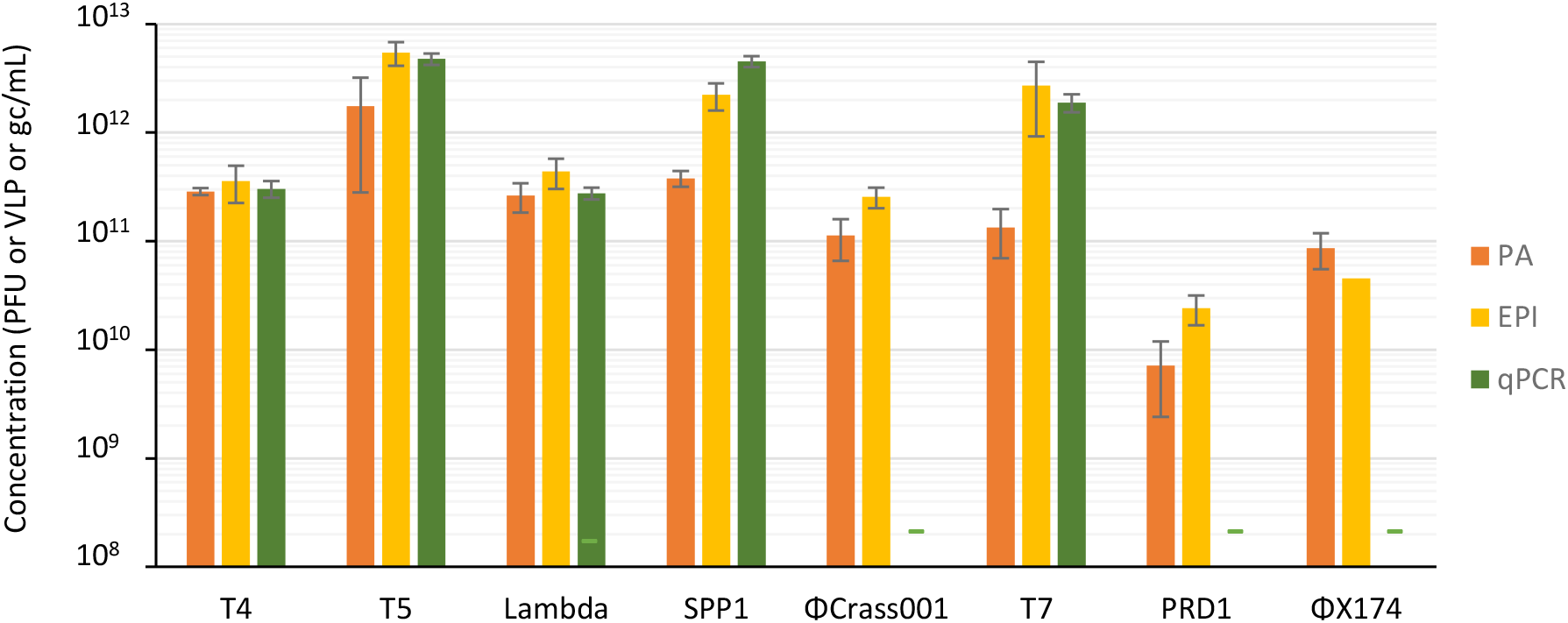
Quantification of phage preparations by standard techniques. Phage concentrations determined by Plaque Assay (PA, yellow), Epifluorescence Microscopy (EPI, orange), and quantitative PCR (qPCR, green). Error bars represent standard error of the mean on three independent experiments. Dashes (-) indicate absence of measurement.

We next compared the performance of the Videodrop and the two NTA devices in determining phage concentrations. First, we measured the concentrations of serial dilutions of T4 phage with the Videodrop (**Fig. 3A**). The optimal concentration range for T4 concentration measurement with the Videodrop matched the range determined previously with the polystyrene beads, i.e., from 3 × 10^8^ to 5 × 10^9^ particles/mL. All further measurements with the Videodrop were therefore carried out in this concentration range. For the NanoSight and Zetaview, we used the concentrations recommended by the manufacturers (material and methods).

**Figure 3.**
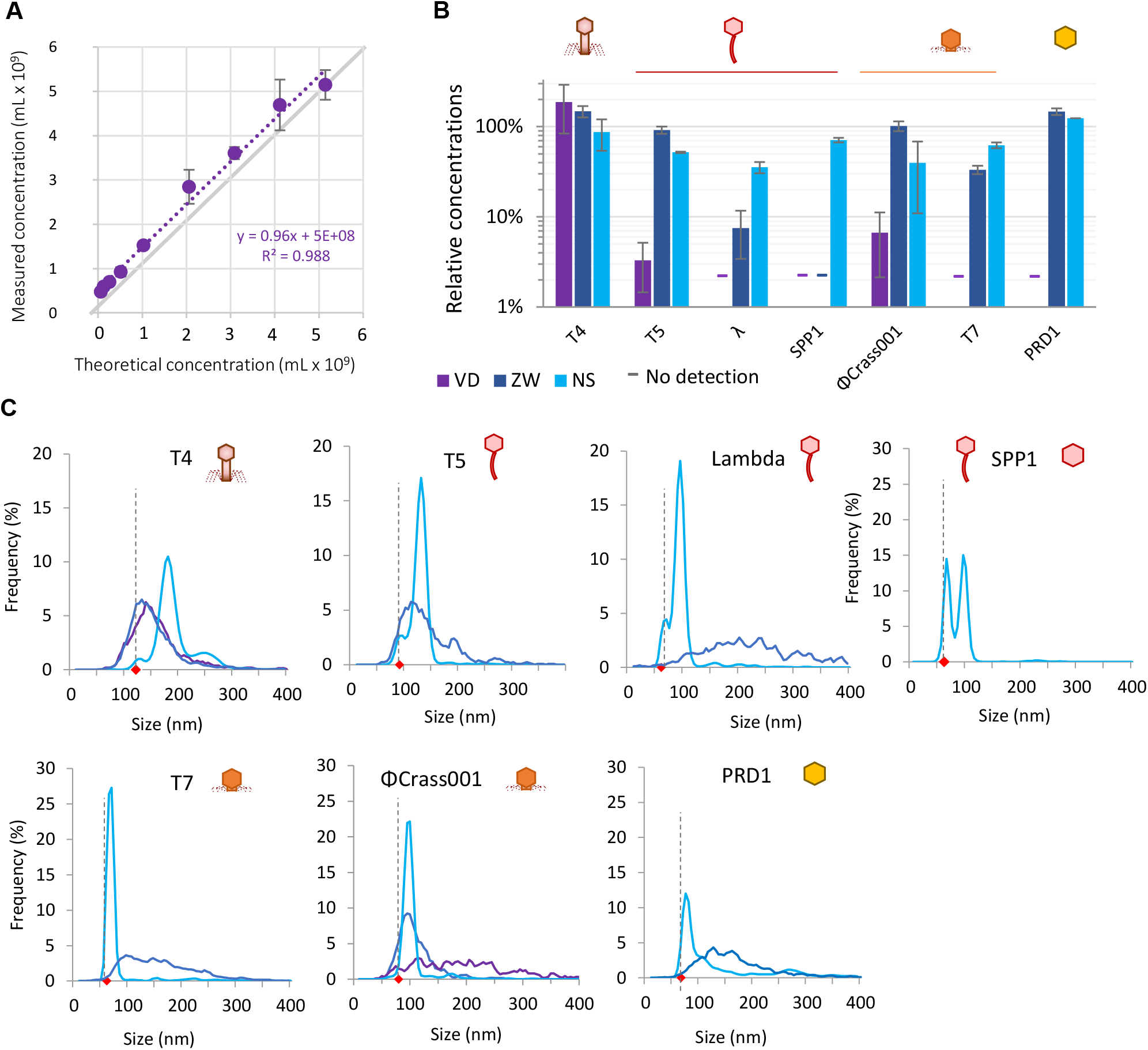
Comparison of ILM and NTA for the analysis of phages. (A) Measurements on serial dilutions of T4 with the Videodrop. (B) Relative phage concentrations obtained with the Videodrop (purple), the ZetaView (dark blue) and the NanoSight (clear blue). Error bars represent standard error of the mean on three independent experiments. Dashes (-) indicate that the device provided no reliable measurement (<1% of EPI values). (C) Virion hydrodynamic diameter (Size) distributions for each device. Colors are as in (B). Black dashed bars and red diamonds indicate the geometrical capsid diameter of phages, as presented in Table 2.

The NanoSight detected all tested phages, except the smallest, ΦX174, for which the capsid diameter is only 30 nm. The NanoSight phage detection threshold was therefore between 30 and 60 nm. The concentrations obtained were in relatively good agreement with the EPI counts, with values that were generally 1-to-3-fold lower for all phages (**Fig. 3B**). The ZetaView also detected most of the phages except two: ΦX174, as the NanoSight, as well as SPP1. This is surprising, as the published diameter of the SPP1 virion, 61 nm, is similar to that of λ and T7 phages (**Table 2**), which were correctly detected. Of note, SPP1 particles create a visible signal that is nonetheless too low to be detected by the ZetaView software with the parameters used in this study, indicating that SPP1 virions are just below the detection threshold. This may result either from a slightly smaller size of the SPP1 virions, as suggested by our TEM observations indicating a diameter of approximately 57 nm (**Table 3**), from a lower refractive index of SPP1 virions relative to other phages, or to smaller surface charge (which results in a smaller hydrodynamic diameter). We therefore considered that the detection threshold of the ZetaView is slightly over 60 nm for phages. For phages with a capsid larger than 60 nm (T4, T5, ΦCrAss001 and PRD1), the ZetaView concentrations were in very good agreement with the EPI counts (**Fig. 3B**).

**Table 3.**
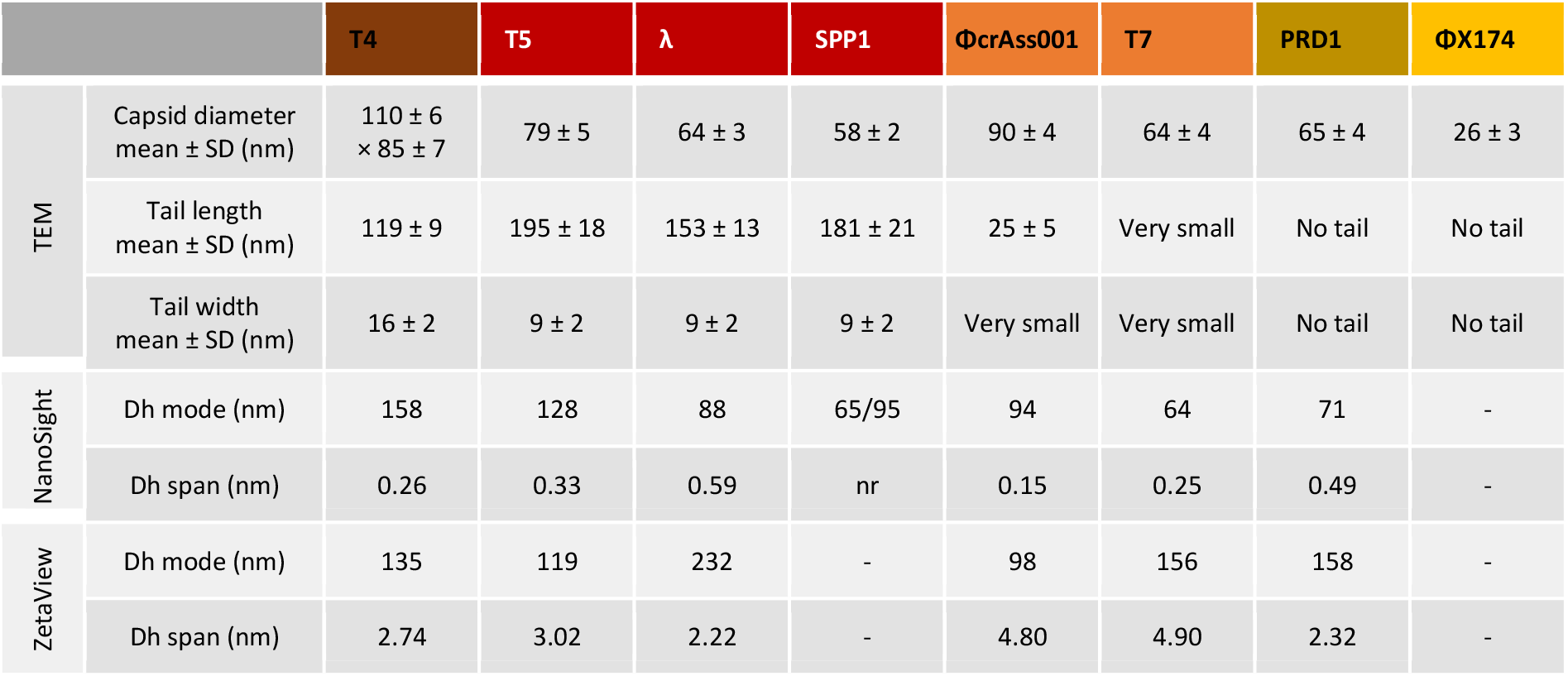
Phage characteristics measured by TEM and NTA. SD : standard deviation, Dh : hydrodynamic diameter, nr: not relevant, -: not measurable. The mode is the value which correspond to the peak of the distribution, and the span = (D90 – D10)/D50, with D10, D50 and D90 being the sizes below which 10%, 50% or 90% of all particles are found respectively. The span gives an indication of how far the 10 percent and 90 percent points are apart, relative to the median.

The Videodrop correctly detected only the largest phage, T4. T5 and ΦCrAss001 virions created visible spots, but the intensity of the signal was too low to be detected by the software, indicating that the particle signal is just below the detection threshold of the device. Given that the size of the T5 and ΦCrAss001 capsid is close to 90 nm (**Table 2**), this suggests that the Videodrop detection threshold is slightly over 90 nm for phages.

In conclusion, the NanoSight and ZetaView correctly enumerated phage particles larger than 50 nm and 60 nm, respectively, consistent with previous studies on synthetic nanoparticles (Bachurski et al. 2019, Dehghani et al. 2021). The Videodrop detected and provided correct concentrations only for T4, a tailed phage with a capsid larger than 90 nm.

### Particle size distributions of phage samples

In addition to the determination of particle concentrations, the three devices provide particle size distributions. The NanoSight provided size distributions in agreement with the samples’ characteristics, i.e., in most cases, narrow unimodal distributions with a maximal value close to the diameter of the virions (**Fig. 3B** and **Table 3**). For the tailless phages (ΦcrAss001, T7 and PRD1), the measured sizes were very close to the geometric diameters determined by microscopy (**Tables 2&3 and Fig. 3**), indicating that the virions are close to perfect spheres. For phages with a large tail, we generally observed a bimodal distribution of sizes, with a first small peak corresponding to the geometric diameter of the capsid, hence to damaged tailless virions, and a second higher peak at higher particle size, most likely corresponding to intact virions, as the presence of a tail is expected to increase the hydrodynamic diameter of the virions (**Fig. 3**). The bimodal distribution was particularly apparent in the case of the SPP1 sample, which was shown by TEM to contain close to 50% tailless capsids (**Fig. 1**).

The ZetaView measured correct modal sizes for T4, T5, and ΦcrAss001, but with poor precision, as evidenced by an important span of the size distributions (**Table 3 and Fig. 3**). Similarly, the Videodrop measured a correct modal size for T4 (133 nm as compared to the geometric diameter of 120 nm), also with poor precision (Fig. 3C). In the case of phages whose diameter is close to the detection threshold (λ, T7 and PRD1), the ZetaView provided particle size distributions that were quite different from the expected distributions in terms of both mode and precision. In conclusion, the NanoSight provided more precise phage size estimations than the other devices, especially on smaller phages.

### Quantification and size distribution of EVs of different origins

We next compared the performance of the three instruments in measuring the concentrations and size distributions of EV preparations from various origins. To this aim, we first prepared EV-enriched preparations (called EVs from here on for simplicity) from nine different origins: bovine milk, THP1 human monocytic cell cultures, *F. prausnitzii, B. subtilis, E. coli*, and *S. aureus* bacterial cultures, and, finally, EVs from germ-free or conventional rodent feces or cecal content. In germ-free animals, EVs exclusively originate from intestinal cells, whereas in conventional animals they also originate from the intestinal microbiota. Observations of EV preparations by TEM confirmed that most of the nanoparticles of the preparations were EVs, with a large dispersion of particle sizes (**Fig. 4**).

**Figure 4.**
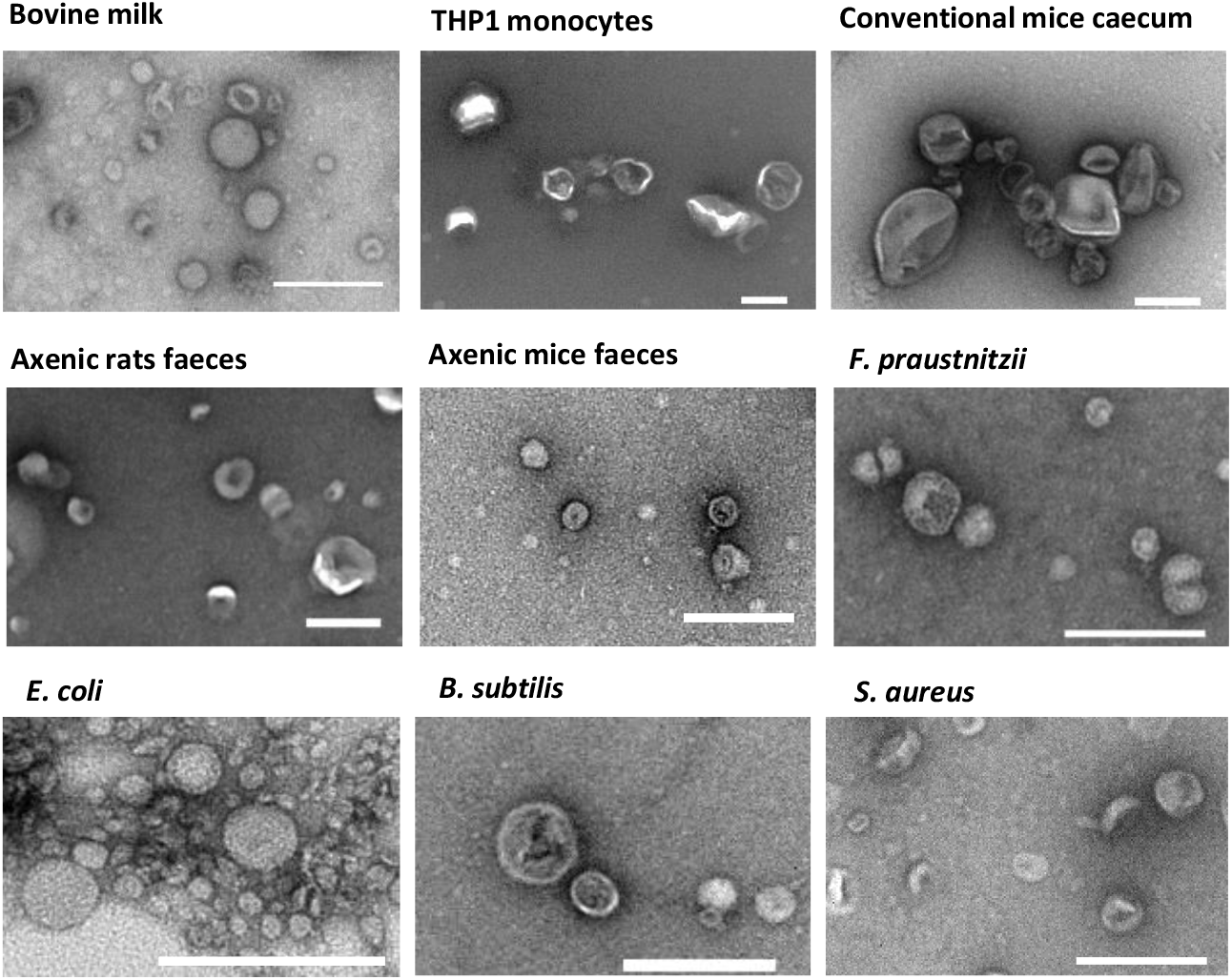
Representative TEM images of the EVs used in the study. Scale bars are 200 nm.

The EV preparations were then examined using the Videodrop, ZetaView, and NanoSight. As optimal concentration ranges vary with the nature of the sample and, in particular, with the level of polydispersity, we first determined the nanoparticle concentrations of serial dilutions of EVs from bovine milk, which range from 35 to 200 nm (**Fig. 5 & 6**). For both NTA instruments, the linearity ranges were much narrower than previously reported for monodisperse samples: they ranged from 2.5 × 10^7^ to only 2 × 10^8^ particles/mL instead of 3 × 10^9^ particles/mL, the upper limit of the linearity range determined with artificial beads ((Maguire et al. 2017). On the contrary, the optimal concentration range for the Videodrop was similar to that determined previously with polystyrene beads and phage particles.

**Figure 5.**
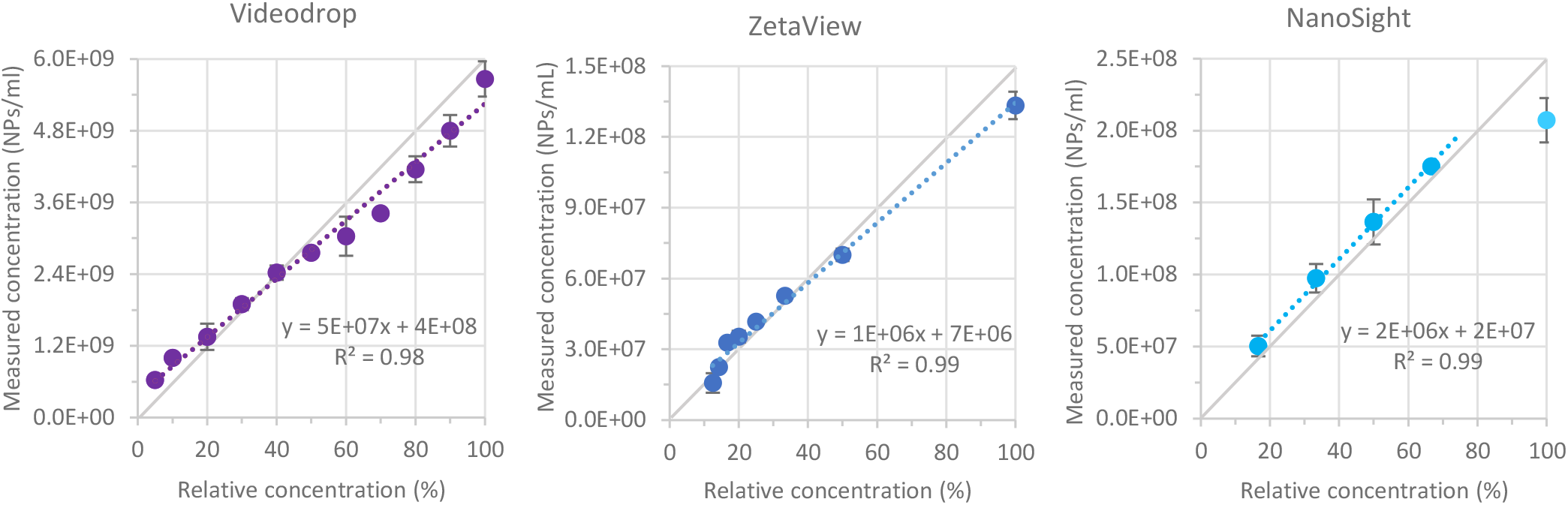
Concentration measurements of serial dilutions of EVs from bovine milk. Successive dilutions were realized in PBS. Error bars represent standard deviation on three independent dilutions.

All EV preparations were then examined with the three optical devices. Contrary to phage samples, the concentrations of EVs were unknown and, therefore, we could only compare the values obtained with the three instruments to each other. Concerning EVs originating at least partially from mammalian cells (i.e., EVs from milk, THP1 monocytes, or mouse intestinal contents), the concentrations determined by the three devices were relatively close, within a two-fold range, the highest concentrations always being provided by the ZetaView (**Fig. 6A**). This is unexpected as TEM suggested the presence, especially in samples from axenic animals, of EVs below the Videodrop detection threshold but larger than those of the NTA devices, even when considering the possible 10 to 30% shrinkage of EVs in TEM (**Fig. 6C**). Of note, the size detection thresholds could not be determined for EVs because of the high polydispersity of the samples, however, given that the refractive index of EVs is only slightly lower than that of phages, the size detection thresholds should be close for both types of objects.

**Figure 6.**
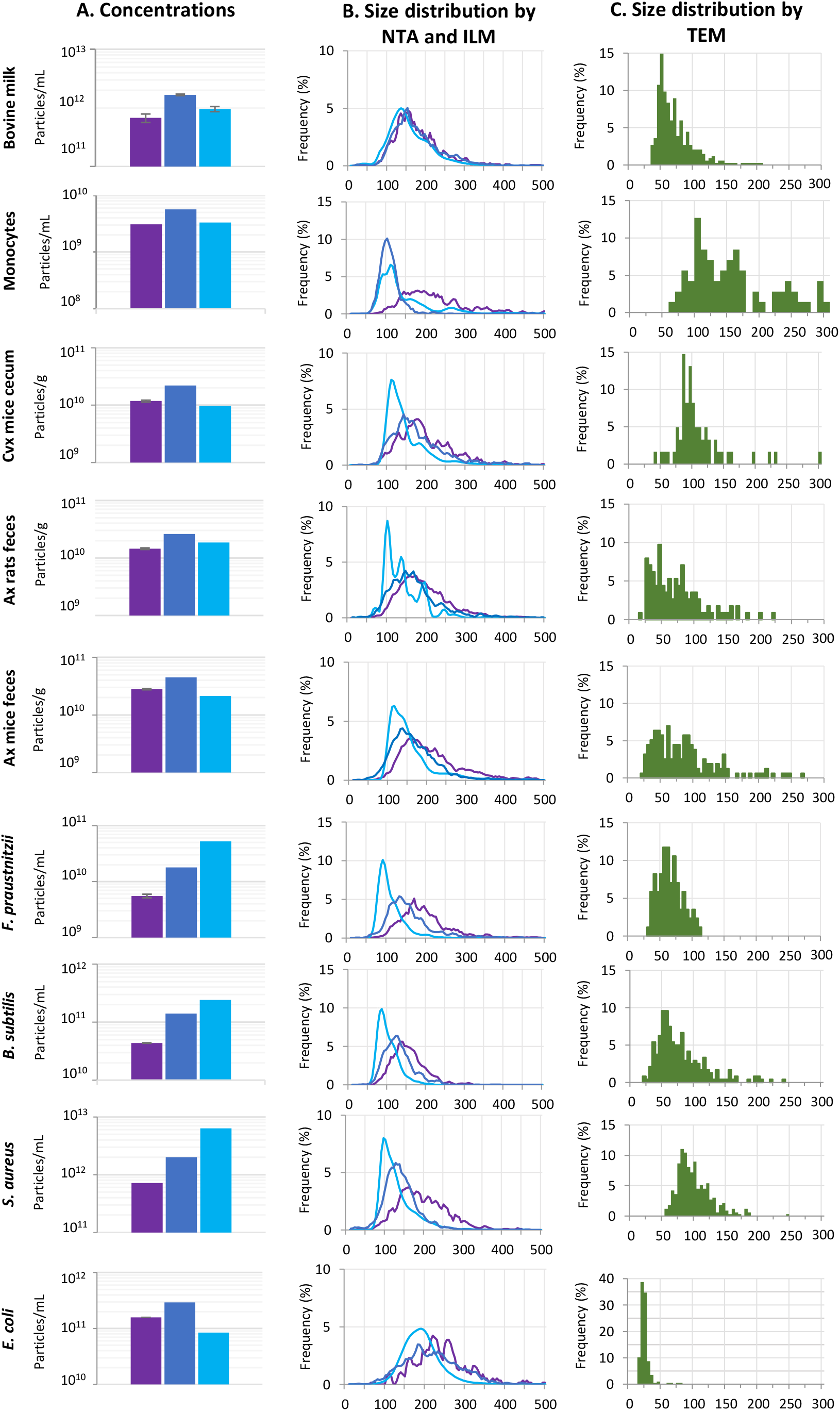
Determination of concentration and size of EVs. (A) Mean concentrations determined by the Videodrop (purple), the ZetaView (dark blue) and the NanoSight (clear blue). (B) Size distributions obtained with the three devices. Colors are as in (A). (C) Size distributions of particles from TEM images on 85 to 350 particles depending on samples. Ax: axenic, Cvx: Conventionnal. Note that the y axis is different in B and C.

By contrast, the devices were discordant in quantifying EVs originating from bacterial cultures, (except for *E. coli*, see below), with up to 10-fold higher concentrations obtained with the NanoSight than with the Videodrop (**Fig. 6B**). The sizes and levels of dispersion observed by TEM being similar in some eukaryotic and prokaryotic EVs (for example compare the size distribution of EVs from axenic mice feces and from *B. subtilis*), this different outcome might results from different characteristics of bacterial and mammalian EVs. The case of *E. coli* EVs is particular, as most EVs had a diameter between 20 and 40 nm, much smaller than the detection thresholds of the three devices. The concentrations measured were therefore probably strongly underestimated with all three devices.

Finally, concerning EV size distributions, all devices provided size distributions that were both shifted toward larger values and broader than the geometrical diameters obtained by TEM (**Fig. 6**). This trend is more pronounced for the Videodrop and ZetaView than for the NanoSight. This suggest that, contrarily to phage virions, EVs are far from being perfect spheres (they are soft and charged) and that hydrodynamic diameters are quite different from geometrical ones.

## DISCUSSION

As a first step towards evaluation of the Videodrop, we precisely determined the concentration of phages using three classical methods. We established that qPCR and EPI provide very similar concentrations, in contrast to a previous study in which the authors found significantly lower concentrations by EPI than qPCR (Kaletta et al. 2020). The better agreement between the values obtained by the two methods in our study may have resulted from the use of a more concentrated staining solution, the addition of a flash freezing step to improve the staining, and/or the use of a different microscopic setup. With our well-characterized phage preparations, we found the size threshold of the Videodrop for phage detection to be approximately 90 nm, as only 5 to 10% of the 80 to 90-nm diameter virions of T5 and ΦcrAss001 were counted. This contrasts with the reported good performance of the Videodrop for the quantification of an adenovirus of 75.5 nm of diameter: in this study, the authors could detect approximately 25% of adenovirus particles (Turkki et al. 2021). Two factors could explain this difference. First, the refractive index of adenoviruses or their surface charge may be higher than that of the phages tested here, improving their detection. Second, the adenovirus observations were carried out using a relative threshold of 3.2, whereas we used a relative threshold of 3.8, consistent with the manufacturer’s recommendations to have few false positive detections.

For the large T4 phage virus (86 × 120 nm), the concentrations provided by the Videodrop instrument were in very good agreement with those obtained by EPI and qPCR. Similar to our observations with polystyrene nanoparticles, a linear count consistent with the dilution was observed for concentrations between 5 × 10^8^ and 4 × 10^9^ particles/mL, indicating the successful enumeration of individual virions. Concerning the NTA instruments, our results suggest a sensitivity threshold for phages < 60 nm of diameter for NanoSight and between 60 and 75 nm for Zetaview under the selected experimental conditions, in accordance with previous results on EVs (van der Pol et al. 2014).

In terms of the determination of nanoparticle size distributions, the NanoSight provided very precise and accurate measurements of capsid sizes for tailless phages, indicating that phage hydrodynamic diameters are close to their geometric diameters. In polydisperse EV samples, however, we show that ILM results in a broader and shifted distribution of EV sizes, similar to what others have already observed for NTA (Bachurski et al. 2019). This difference is too great to be solely explained by the shrinking phenomenon sometimes observed with TEM (Chernyshev et al. 2015, Kotrbova et al. 2019). Instead, this overestimation most likely results from several phenomena. First and probably most importantly, EVs smaller than the detection threshold of the devices (60 to 70 nm for NTA and 90 nm for ILM) are not detected by NTA. Small particles close to the detection limit are difficult to track, due to low intensity and rapid movement, and therefore often cannot be attributed a size. Second, the protein surface cargo influences the hydrodynamic diameter measured by NTA (Skliar et al. 2018, Bachurski et al. 2019), resulting in sizes measured by NTA larger than the geometric diameters. Finally, in NTA-based devices, large particles mask small ones (Ortega Arroyo et al. 2016, Dehghani et al. 2021). In theory, ILM is less subject to such a masking effect because the detected signal is proportional to the third power of the diameter of the nanoparticles, whereas in NTA, the detected signal is proportional to the sixth power of the particle’s diameter. This could explain why, despite the lower sensitivity of the Videodrop instrument, it was able to determine EV concentrations similar to those of the NTA instruments in milk and mouse intestinal samples, which are highly polydisperse in size.

In any case, size measurements provided by NTA and ILM instruments must be considered with caution, as they do not reflect the true geometrical size distribution. The case of *E. coli* EVs particularly highlights the importance of coupling TEM observations with ILM or NTA. Indeed, in this sample, TEM showed that most vesicles were < 50 nm in diameter, smaller than the detection threshold of ILM and NTA. Thus, the concentrations obtained with these devices were clearly underestimations.

On another level, a major advantage of the Videodrop instrument is its ease and speed of use. In particular, sample positioning is very rapid (a 6-μL drop of the sample has to be deposited into a dedicated microcavity) relative to positioning of the sample through a syringe in NTA, which has to be very carefully connected to the tubing to avoid generating air bubbles, increasing the required time of manipulation (**Table 4**). The ZetaView presents similar limitations as the NanoSight; however, although it necessitates a calibration step prior use, this device offers better efficiency in terms of the time of analysis. The ultra-fast measurements possible with the Videodrop is of particular importance when a large number of samples have to be measured. Another advantage of the sample positioning of the Videodrop is that it enables the measurement of samples rich in diverse contaminants, such as soluble lipids and proteins, which is not possible with the NTA devices, as such contaminants could result in clogging of the tubing. For example, in the context of this study, raw fecal filtrates could not be observed with the NTA instruments, as they contain flagella, lipids, and proteins that are either soluble or in aggregates, whereas such raw filtrates could be observed with the Videodrop.

**Table 4.**
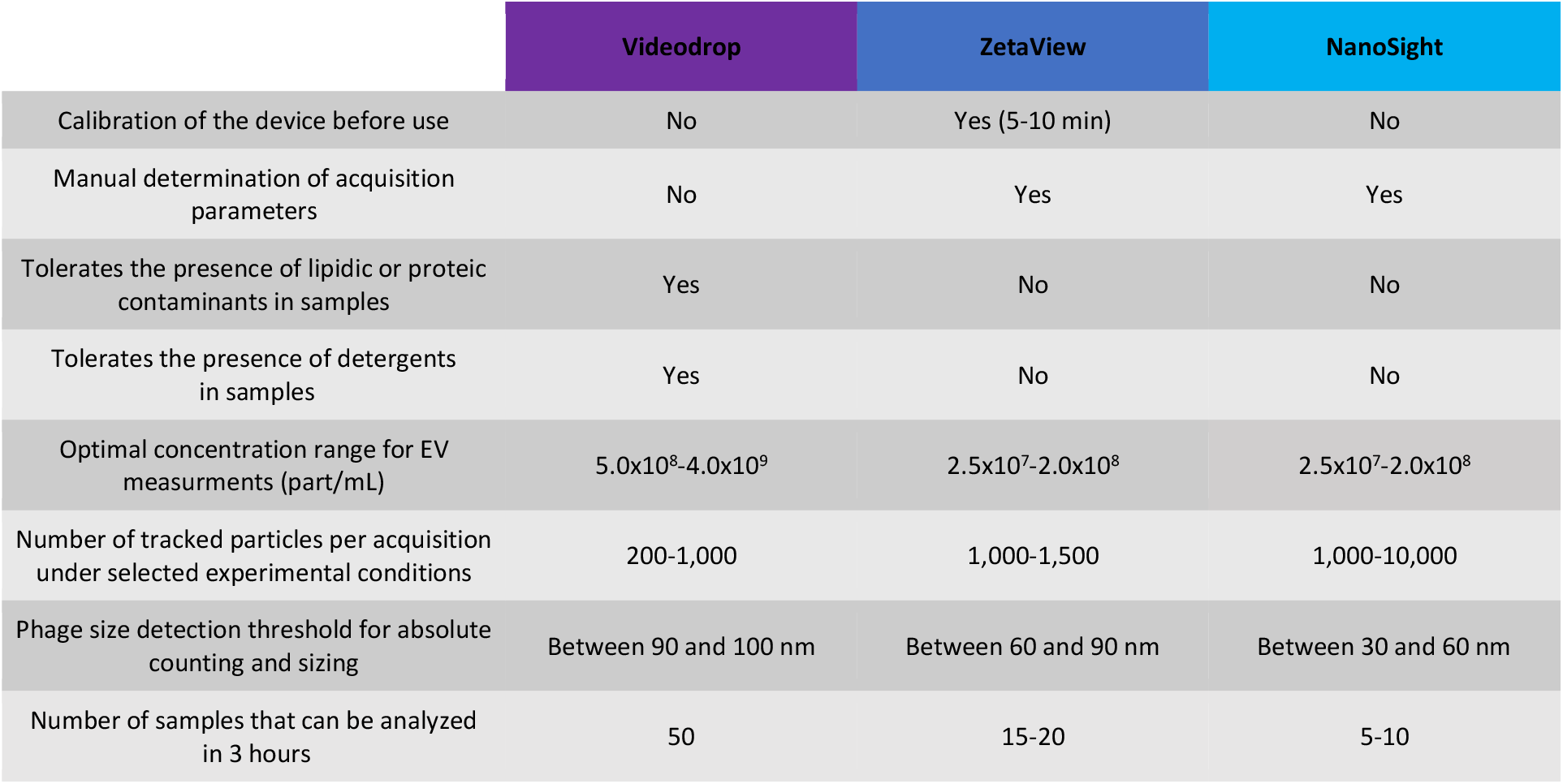
Evaluation summary of the Videodrop, ZetaView and NanoSight

A final advantage of the Videodrop is the automatic adjustment of the acquisition parameter for the video capture, here LED intensity. By contrast, NTA requires several optimization steps to determine the suitable settings for video capture and analysis. As reported previously (Maas et al. 2015, Bachurski et al. 2019), the camera settings in both devices (NanoSight NS300: camera level, ZetaView: camera sensitivity) have a profound impact on the measured concentration. In addition, the optimal settings are very difficult, if not impossible, to determine for polydisperse samples, as raising the level of the camera results in the better observation of small particles but also increased noise created by large ones (and, conversely, lowering the level of the camera is better for large-particle enumeration but leads to the loss of small particles).

In conclusion, the better sensitivity of the NanoSight instrument makes it the most appropriate device for concentration determinations of monodisperse populations of objects between 50 and 90 nm in diameter. In addition, of the three devices we tested, the NanoSight provided the most precise size distributions and projects requiring precision on a limited number of samples would benefit the most from this device. However, the higher precision and sensitivity of the NanoSight is offset by four disadvantages: its time-consuming manipulation, its high susceptibility to masking effects, the impossibility to analyze samples that contain a large number of impurities, and its price. Depending on the application, the most important advantage of the Videodrop would be its ease of use and the rapidity of sample observation (**Table 2**). In addition, ILM is less prone to small-particle masking by larger particles and thus compares favorably with NTA for concentration measurements of highly polydisperse populations. Finally, the Videodrop is less expensive and requires fewer consumables than the other devices (a simple pipet tip as opposed to a syringe), which is directly related to a lower environmental footprint, which should also be taken into consideration.

## ACKNOWLEDGEMENTS

The authors thank Christine Longin from the MIMA2 Micalis facility and Julien Lossouarn for their help with the acquisition of the TEM images and Paulo Tavares for providing the SPP1 phage.

