## supplementary figures for "Comparison of interferometric light microscopy with nanoparticle tracking analysis for the study of extracellular vesicles and bacteriophages"

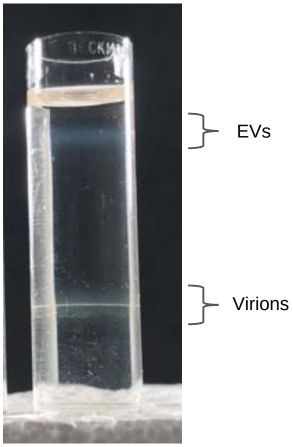


**Figure S1**. **Picture of a representative iodixanol gradient.** The upper band is at the density expected for EVs, while the lower one is at the density of most phages.

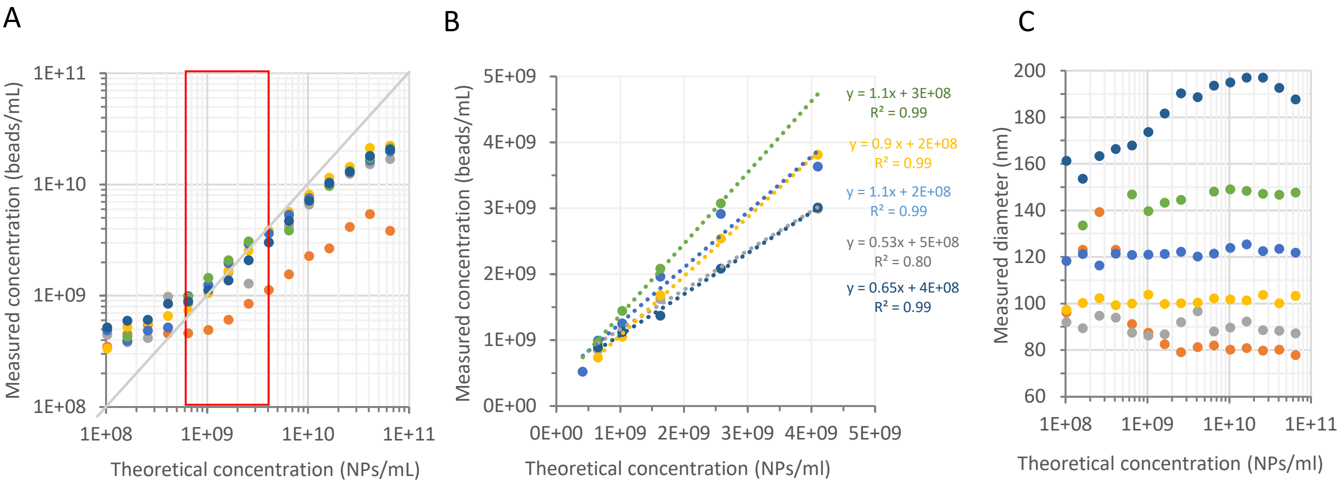


**Figure S2**. **Concentrations and sizes of serial dilutions of polystyrene beads.**

(A) Measured concentrations in function of theoretical concentrations from $10^{11}$ to $10^{8}$ beads per mL. The red rectangle delimits the concentration range in which a linear count consistent with the dilution was observed ($5x10^{8}$ to $4x10^{9}$ beads/mL). (B) Measured concentrations in function of theoretical concentrations between $5x10^{8}$ to $4x10^{9}$ beads per mL. Equations and coefficient of determination of linear fits are indicated for each bead size. (C) Measured diameters of beads as a function of their concentration.

| Phage | Primers | Reference |
|---|---|---|
| **T4** | F: ACTGGCCAGGTATTCGCA<br>R: ATGCTTCTTTAGCACCGGCA | (Klopot 2017) |
| **T5** | F:AGCCCGTGAGAAACTGTACA<br>R:CTGCTCCAGTATCCGTCAG | This article |
| **λ** | F: GAGTGCGGAAGATGCAAAGG<br>R: TTAACAGTGCGTGACCAGG | (Fogg 2010) |
| **SPP1** | F: CGGGCTGAAATACCTGTGGA<br>R: TAGCCCCTCCTCCGATTGTT | (Labarde 2021) |
| **T7** | F: CCCGAACTATAAGGCTAACC<br>R: GCTCACGGATGCAATAGAAC | This article |

**Supplementary Table 1. qPCR primers used in this study**
